# Histone Variant H3.3 Mediates DNA Repair and cGAS-STING Pathway Activation in Telomere Dysfunction

**DOI:** 10.1101/2024.08.07.606966

**Authors:** Chiao-Ming Huang, Liuh-Yow Chen

## Abstract

The telomere damage response is a critical mechanism regulating human aging and disease progression. Deprotected telomeres activate the cytosolic DNA-sensing cGAS-STING pathway; however, the underlying mechanism remains unclear. Here, we discover that histone H3.3 is required for cGAS-STING pathway activation in cells with deprotected telomeres. Expression of the TRF2 dominant-negative mutant, TRF2ΔBΔM, induces telomere dysfunction in fibroblast cells, triggering cGAS-STING pathway activation and growth inhibition. Histone H3.3 depletion significantly reduces this activation, highlighting its critical role in linking telomere deprotection to the cGAS-STING-mediated innate immune signaling. Furthermore, we assess the role of histone H3.3 in telomere fusion. Our finding reveal that histone H3.3 regulates cGAS-STING signaling by controlling telomere fusion. Additionally, depletion of histone H3.3 chaperones, including ATRX, DAXX, and HIRA, inhibits telomere fusion and cGAS-STING pathway activation, underscoring the role of histone H3.3 in telomere maintenance and the DNA damage response. Collectively, our study establishes histone H3.3 as a key regulator of telomere fusion and telomere dysfunction-induced cGAS-STING pathway activation, revealing the correlation between histones and telomeric DNA damage response.

## INTRODUCTION

Telomeres and their associated shelterin proteins are crucial for protecting chromosome ends^1^. Deprotection of telomeres triggers a DNA damage response (DDR), leading to replicative senescence^2–4^. Among the sheltering proteins, telomeric repeat binding factor 2 (TERF2, TRF2) plays a critical role in preventing DDR at chromosome ends. Depletion of TRF2 from telomeres induces cellular senescence and chromosome end-to-end fusion^3^, an event that is often found when telomeres are deprotected, particularly under conditions involving dominant-negative TRF2 expression^3^. Previous studies demonstrated that the absence of functional TRF2 activates a DDR mediated by the ataxia telangiectasia mutated (ATM) pathway^3–5^. Upstream components of ATM, including the double-strand break (DSB) sensing complex, the meiotic recombination 11 (MRE11)-radiation sensitive 50 (RAD50)-Nijmegen breakage syndrome 1 (NBS1, NBN) (MRN) complex, play a crucial role in sensing TRF2-dependent telomere deprotection^6–8^. Conversely, downstream effector proteins of ATM, such as Ku protein 70/80 (Ku70/80), ligase IV (LIG4), and x-ray repair cross-complementing 4 (XRCC4), mediate the signaling pathways induced by TRF2 depletion and subsequent telomere fusion^3,9,10^. Together, these findings establish an ATM-dependent mechanism underlying telomere deprotection and the induction of cellular senescence.

Beyond ATM-dependent mechanisms, the cGAS-STING pathway is implicated in telomere dysfunction^11^. In this pathway, cyclic GMP-AMP synthase (cGAS) and stimulator of interferon response cGAMP interactor (STING) collaborate to activate a phosphorylation cascade involving TANK-binding kinase 1 (TBK1), STING, and interferon regulatory factor 3 (IRF3), which drives the innate immune response by promoting type I IFN transcription^12^. Studies demonstrate that telomere-damaging agents and defective telomere-binding proteins can lead to micronuclei formation, thereby activating the cGAS-STING pathway^11^. Furthermore, evidence supports the involvement of the cGAS-STING pathway in telomere deprotection-induced cellular senescence. For example, chromatin bridges formed during dominant-negative TRF2 expression are processed by three prime repair exonuclease 1 (TREX1)^13^, resulting in DNA fragments that activate the cGAS-STING pathway^14^. Similarly, the mutated TRF2, TRF2ΔB induces the formation of extrachromosomal telomere repeats (ECTRs), which are released into the cytosol to trigger the cGAS-STING-mediated innate immune response^15^. Additionally, late-generation mice with deletion of the telomerase RNA component Terc also display senescent signatures along with aberrant STING activation^16^. These findings suggest that telomere deprotection activates both ATM dependent and cGAS-STING pathway, contributing to cellular senescence.

At telomeres, the specific histone variant, histone H3.3, is enriched and plays a crucial role in telomere maintenance^17–19^. Histone H3.3 is deposited in a replication-independent manner by the histone cell cycle regulator (HIRA) complex on euchromatin and by the alpha-thalassemia/mental retardation syndrome X-linked (ATRX)-death domain associated protein (DAXX) complex on heterochromatin, centromeres, and telomeres^18^. While histone H3.3 is known to regulate the active expression of genes^20–24^, its function at telomeres remains unclear. Mutations in histone H3.3, such as histone H3.3^K27M^ and histone H3.3^G34V^, are associated with glioblastoma and other telomerase-negative cancers that utilize the alternative lengthening of telomeres (ALT) pathway^25–28^. ALT, a recombination-based telomere elongation mechanism, is characterized by reduced ATRX expression^28–33^ and impaired cGAS-STING pathway activation^15^. Given the role of ATRX-DAXX in depositing histone H3.3 at telomeres, mutations in ATRX, DAXX, and histone H3.3 may alter telomere dynamics and contribute to ALT-associated cancer development^29,34–36^. These observations underscore histone H3.3’s potential roles in regulating telomere maintenance.

Previously, we reported that depletion of histone H3.3, ATRX, and DAXX impairs the cGAS-STING pathway activated by ECTRs^15^. Our findings indicated that histone H3.3 participated in the telomere-associated cGAS-STING pathway activation. Additionally, histone H3.3 is involved in DNA damage repair^37^, yet the relationship of histone H3.3’s roles in regulating DNA damage responses at telomeres and activating the cGAS-STING pathway remains unclear. Here, we demonstrated a role of histone H3.3 in telomere deprotection, the activation of the cGAS-STING pathway, and deprotected telomere-induced cell growth inhibition.

## RESULTS

### TRF2ΔBΔM expression drives cGAS-mediated growth inhibition

To investigate the signaling pathway induced by the telomeric deprotection, we focused on exploring the downstream cellular response in cells lacking TRF2. We developed an inducible expression system using a Tet-On mechanism to express dominant-negative mutant TRF2ΔBΔM in telomerase-immortalized BJ human fibroblasts (BJ^t^), under the control of a tetracycline-dependent promoter. TRF2ΔBΔM, a deletion mutant of TRF2 lacking its N-terminal basic domain and C-terminal DNA binding domain, disrupts the formation of a functional telomere shelterin complex by displacing endogenous TRF2, leading to telomere deprotection and fusion^3^. Upon adding doxycycline (DOX), we observed the expression of myc-tagged TRF2ΔBΔM (Fig. 1A) and the formation of telomere dysfunction-induced foci (TIFs), characterized by the colocalization of phosphorylated histone H2AX (γH2AX) with telomeres (Fig. 1B) in BJ^t^ cells.

**Figure 1.**
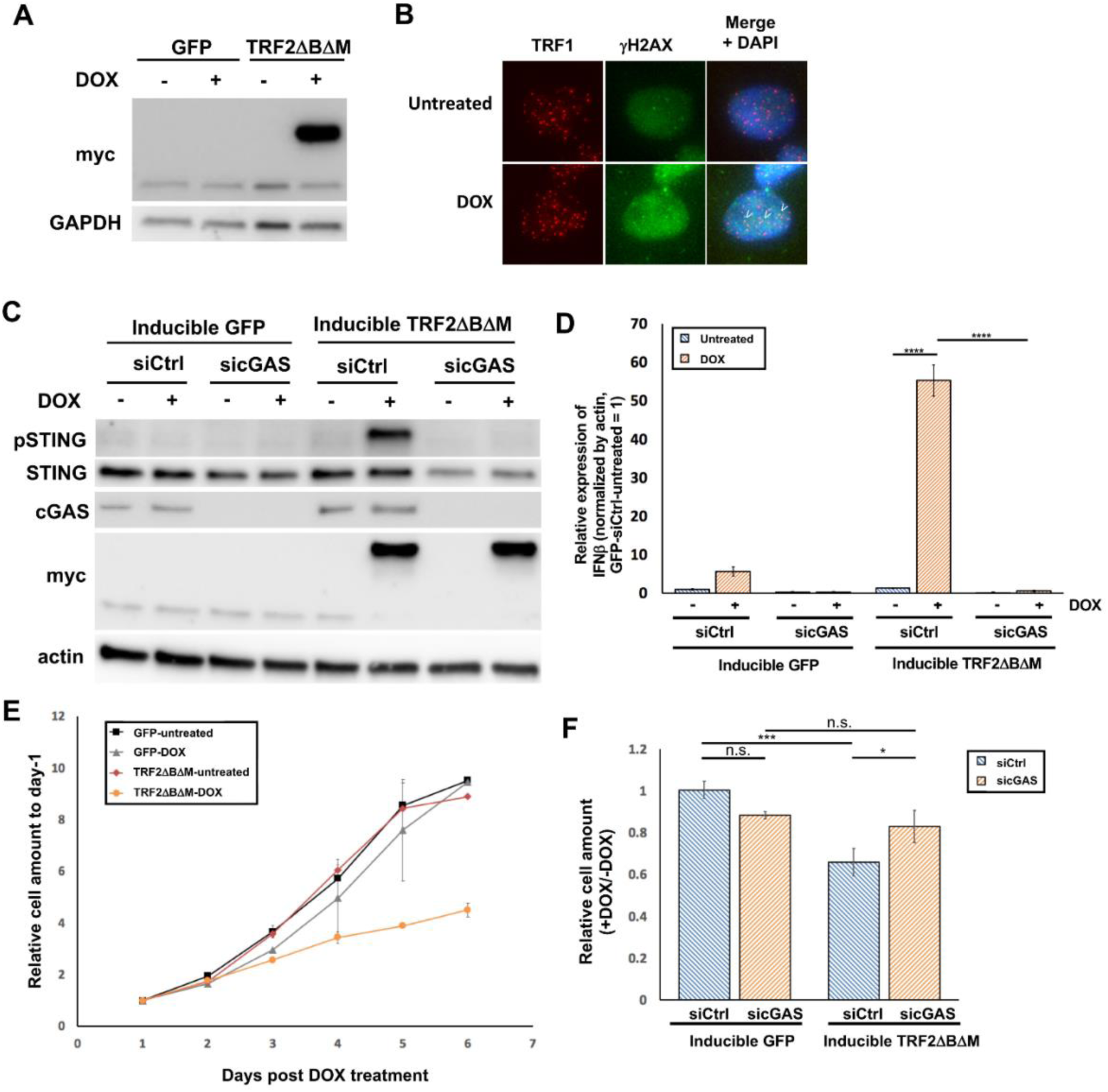
Expression of TRF2ΔBΔM triggers cGAS-STING pathway activation and cGAS-mediated growth inhibition. BJ^t^ cells were separated into untreated and treatment groups, and then 25 ng/ml doxycycline (DOX) was mixed into the culture medium of the treatment group to induce expression of TRF2ΔBΔM. (**A**) Ectopic expression of myc-tagged TRF2ΔBΔM and GAPDH and was determined by Western blotting. (**B**) Immunostaining for γH2AX and TRF1 was performed on day 4 after DOX treatment. Representative images stained for γH2AX (green) and TRF1 (red). White arrowheads indicate representative colocalized signals of TRF1 and γH2AX. For silencing the expression of cGAS, siRNAs were added twice, i.e., at 60 h and 12 h before inducing TRF2ΔBΔM. (**C**) Protein levels of STING phosphorylated at residue S366 (pSTING), STING, cGAS, myc, and actin were determined by Western blotting. (**D**) The expression level of IFNβ mRNA was determined by RT-qPCR. qPCR experiments represent the mean ±SD of three technical repeats. (**E**) Numbers of cells surviving under the indicated conditions were determined by counting the DAPI signals in the culture dishes. Cell numbers were normalized to day 1 after treatment for each experiment. Data represent the mean ±SD of three biological repeats. (**F**) BJ^t^ cells inducibly expressing GFP or TRF2ΔBΔM and respective controls were treated with or without 25 ng/ml DOX. Surviving cells were counted 4 days later. Statistical analysis was conducted using one-way ANOVA. n.s. represents no significant differences, * p-value < 0.05, *** p-value <0.001, and **** p-value < 0.0001.

In line with our aims, we observed the activation of cGAS-STING response following TRF2ΔBΔM induction. Notably, STING phosphorylation, a critical event for the cGAS-STING pathway that facilitates the phosphorylation of IRF3 to drive IFN expression^38,39^, was detected on day 2 and peaked on day 3 post-DOX treatment, indicating that cGAS-STING pathway was activated (Fig. S1A). This activation correlated with the induction of TRF2ΔBΔM expression (Fig. S1A). Reverse-transcription quantitative polymerase chain reaction (RT-qPCR) revealed upregulated expression of IFN beta (IFNβ) mRNA after induced expression of TRF2ΔBΔM (Fig. S1B). These findings suggest that induced expression of TRF2ΔBΔM drives downstream responses including the STING phosphorylation and the transcription of IFNβ.

To determine whether cGAS plays a role in STING activation upon TRF2ΔBΔM expression, we depleted it using small interfering RNAs (siRNAs) before DOX induction. Depletion of cGAS inhibited STING phosphorylation and IFNβ transcription in the TRF2ΔBΔM-expressing cells (Fig. 1C, D), supporting the conclusion that TRF2ΔBΔM activates the cGAS-STING signaling cascade to promote IFNβ production. These results are reminiscent of those from previous studies indicating that TRF2 inhibition activates the cGAS-STING pathway^11,15^.

Furthermore, previous reports have described telomere deprotection induced by TRF2ΔBΔM expression as a trigger for cell growth arrest^3^. Consistently, we also observed that DOX-induced TRF2ΔBΔM expression in BJ^t^ cells led to significantly reduced cell growth (Fig. 1E). Since cGAS-STING pathway activation induces IFNβ expression and affects cell proliferation^15^, we hypothesized that the cell growth defects associated with TRF2ΔBΔM expression might be driven by cGAS-STING pathway activation. To test this possibility, we transfected BJ^t^ cells with siRNAs to silence cGAS expression and monitored their subsequent cell growth. Our results demonstrated that cGAS depletion partially rescued the cell proliferation defects of BJ^t^ cells expressing TRF2ΔBΔM (Fig. 1F), indicating that induction of the cGAS-STING pathway contributes to TRF2ΔBΔM-mediated cell growth defects. Thus, the cGAS-STING pathway exerts a fundamental role in mediating cellular growth impairment due to telomere deprotection.

### Histone H3.3 regulates the cGAS-STING-pathway-mediated cell growth inhibition upon telomere dysfunction

Previously, we showed that histone H3.3 and its chaperone, the ATRX-DAXX complex, promote the activation of cGAS-STING pathway^15^. This activation occurs in response to the accumulation of ECTRs following artificially induced telomere excision by another TRF2 mutant, TRF2ΔB, which lacks only the N-terminal basic domain^15^. This finding demonstrates that the release of telomeric DNA into the cytosol, a phenomenon commonly observed in ALT cancers^40–42^, activates cGAS-STING pathway. Given that ectopic expression of either TRF2ΔB or TRF2ΔBΔM elicits TRF2 dysfunction, we sought to investigate if histone H3.3 is involved in the cGAS-STING pathway activation driven by TRF2ΔBΔM-induced telomere deprotection. To do so, we used siRNAs to deplete histone H3.3 from BJ^t^ cells and then examined TRF2ΔBΔM-induced STING phosphorylation and IFNβ production. Our results revealed that histone H3.3 depletion significantly reduced STING phosphorylation and IFNβ expression in BJ^t^ cells expressing TRF2ΔBΔM (Fig. 2A, B). Furthermore, siRNA-mediated histone H3.3 silencing alleviated the phenotype of defective cell proliferation associated with TRF2ΔBΔM (Fig. 2C). Importantly, the phosphorylation of STING can be rescued by ectopically expressing the siRNA-insensitive H3.3 (Fig. 2D). Collectively, our findings suggest that histone H3.3 mediates cGAS-STING pathway activation and the subsequent cell growth impairment arising from telomere deprotection.

**Figure 2.**
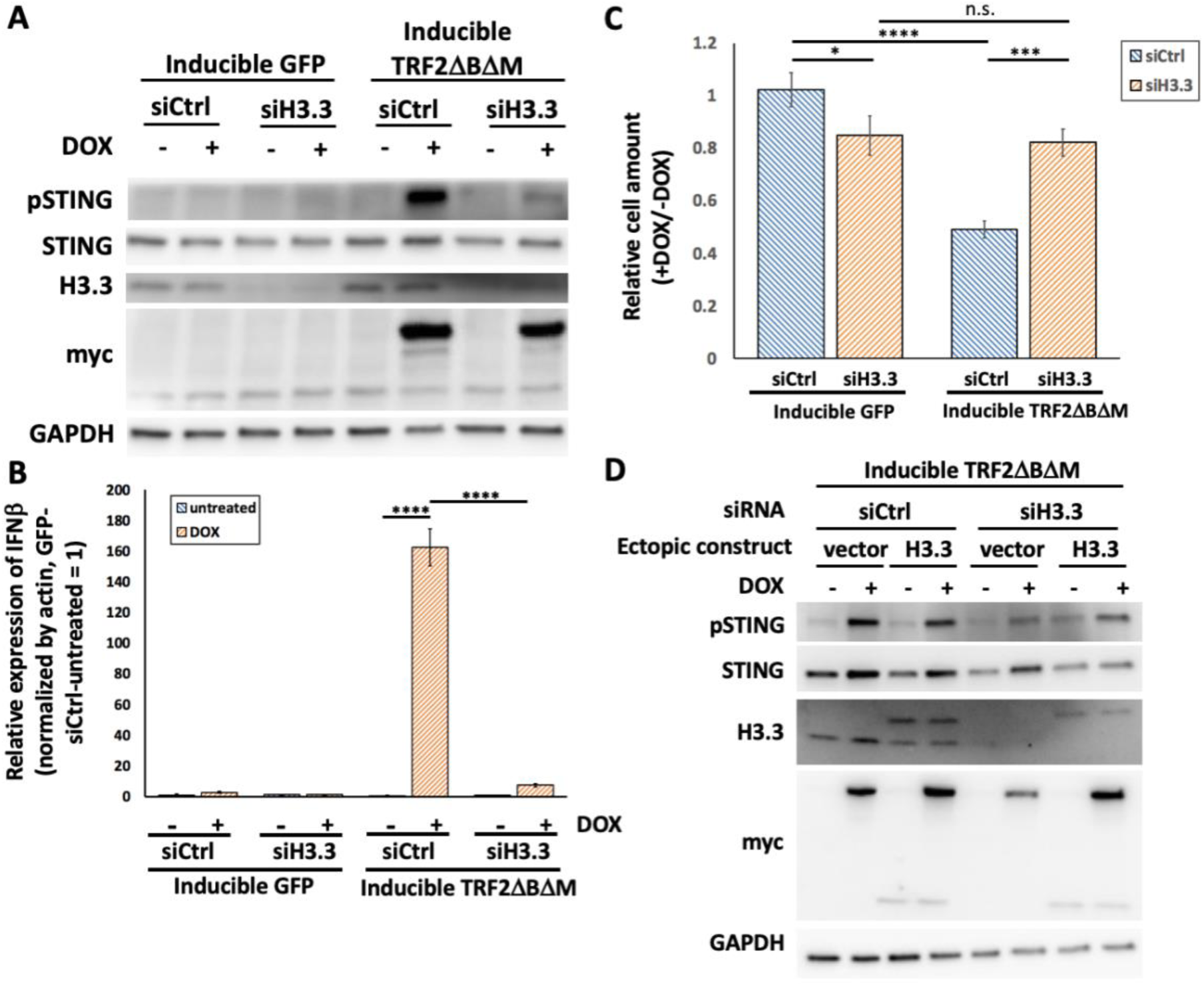
Depletion of H3.3 impairs cGAS-STING pathway activation and rescues growth defects due to the expression of TRF2ΔBΔM. (**A**) BJ^t^ cells stably expressing GFP or myc-tagged TRF2ΔBΔM were established. siRNAs were then added at two different times, i.e., at 60 h and 12 h before inducing TRF2ΔBΔM expression through doxycycline (DOX). Samples were cultured for 2.5 days to investigate the cGAS-STING signaling response. Protein levels of phosphorylated STING (pSTING), STING, H3.3, myc, and GAPDH were determined by Western blotting. (**B**) mRNA expression levels of IFNβ were determined by RT-qPCR. qPCR experiments represent the mean ±SD of three technical repeats. (**C**) Numbers of cells in the DOX+ or DOX-treatment groups were counted to evaluate cell growth 4 days later. Data represent the mean ±SD of three biological repeats. (**D**) A siRNA-insensitive H3.3 construct was introduced into cells via viral vectors and ectopically expressed. Protein levels of phosphorylated STING (pSTING), STING, H3.3, myc, and GAPDH were then determined by Western blotting. Statistical analysis was conducted using one-way ANOVA. n.s. represents no significant differences, * p-value < 0.05, *** p-value <0.001, and **** p-value < 0.0001.

### Telomere fusion by non-homologous end joining (NHEJ) promotes cGAS-STING pathway activation

Telomere deprotection resulting from TRF2 deficiency leads to telomere fusion via the NHEJ pathway^3,10,43^, which in turn generates micronuclei^11,44^ and activates the cGAS-STING pathway^11^. While some studies debate whether micronucleus formation directly triggers cGAS-STING pathway activation^45^, we hypothesized that cGAS-STING pathway activation following TRF2ΔBΔM expression may result from telomere fusion. To test this hypothesis, we used siRNAs to silence the expression of the LIG4 and XRCC4, two key proteins that mediate telomere fusion via the NHEJ mechanism^10^. We found that depletion of LIG4 or XRCC4 in BJ^t^ cells reduced TRF2ΔBΔM-induced telomere fusion, as evidenced by the diminished apparent convergence of telomere signals at chromosome ends (Fig. 3A, B). Furthermore, silencing LIG4 or XRCC4 decreased the formation of abnormal nuclear structures, including nucleoplasmic bridges, nuclear buds, and micronuclei (Fig. 3C, D). Notably, LIG4 or XRCC4 depletion also inhibited STING phosphorylation and IFNβ mRNA induction in BJ^t^ cells expressing TRF2ΔBΔM (Fig. 4). These results suggest that activation of the cGAS-STING pathway in response to TRF2ΔBΔM expression may be attributable to NHEJ-mediated fusion of deprotected telomeres.

**Figure 3.**
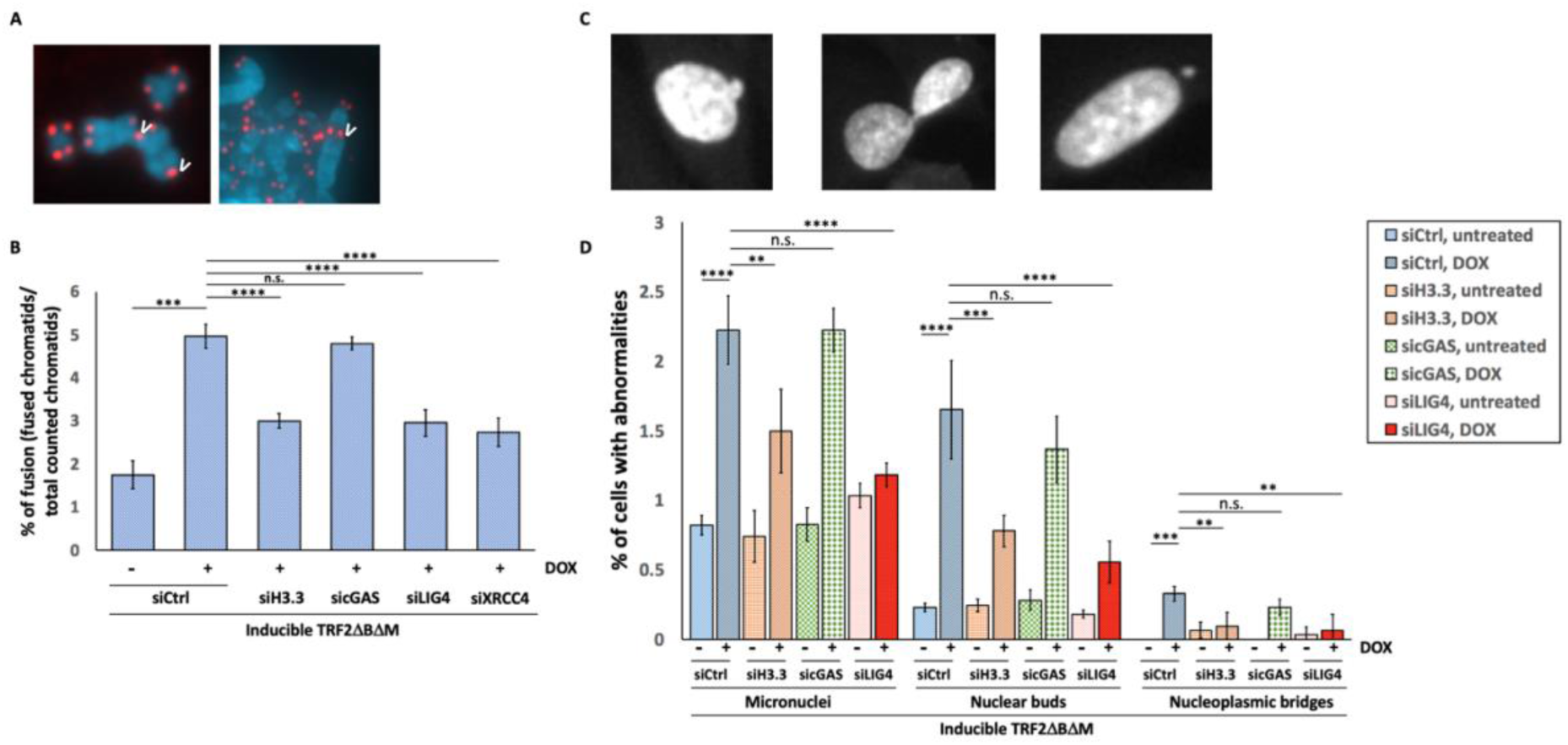
Depletion of H3.3 expression limits telomere-to-telomere fusion events and the formation of nuclear abnormalities. BJ^t^ cells were treated with siRNAs including siCtrl, siH3.3, sicGAS, siLIG4, and siXRCC4 at 60 h and 12 h before DOX treatment to induce expression of TRF2ΔBΔM for 24 h. Cells were arrested using 0.1 μg/ml Demecolcine for another 24 h, and then spread on a slide to visualize telomere-to-telomere fusion events by staining with TelC probes. (**A**) Representative images of telomere fusion events (white arrows). (**B**) Quantitative results for telomere fusion events were calculated from >2000 chromosomes for each sample. Data represent mean ±SD of three biological repeats. (**C**) Representative cells images of siRNA-treated cells on day 2 after DOX treatment. For each group, >1,000 nuclear images were acquired and counted to determine percentages of the indicated nuclear abnormalities. Representative images are shown for a nuclear bud (left), a nucleoplasmic bridge (middle), and a micronucleus (right). (**D**) Quantitative results for nuclear abnormalities. Data represent the mean ±SD of three technical repeats. Statistical analysis was conducted using one-way ANOVA. n.s. represents no significant differences, ** p-value < 0.01, *** p-value <0.001, and **** p-value < 0.0001.

**Figure 4.**
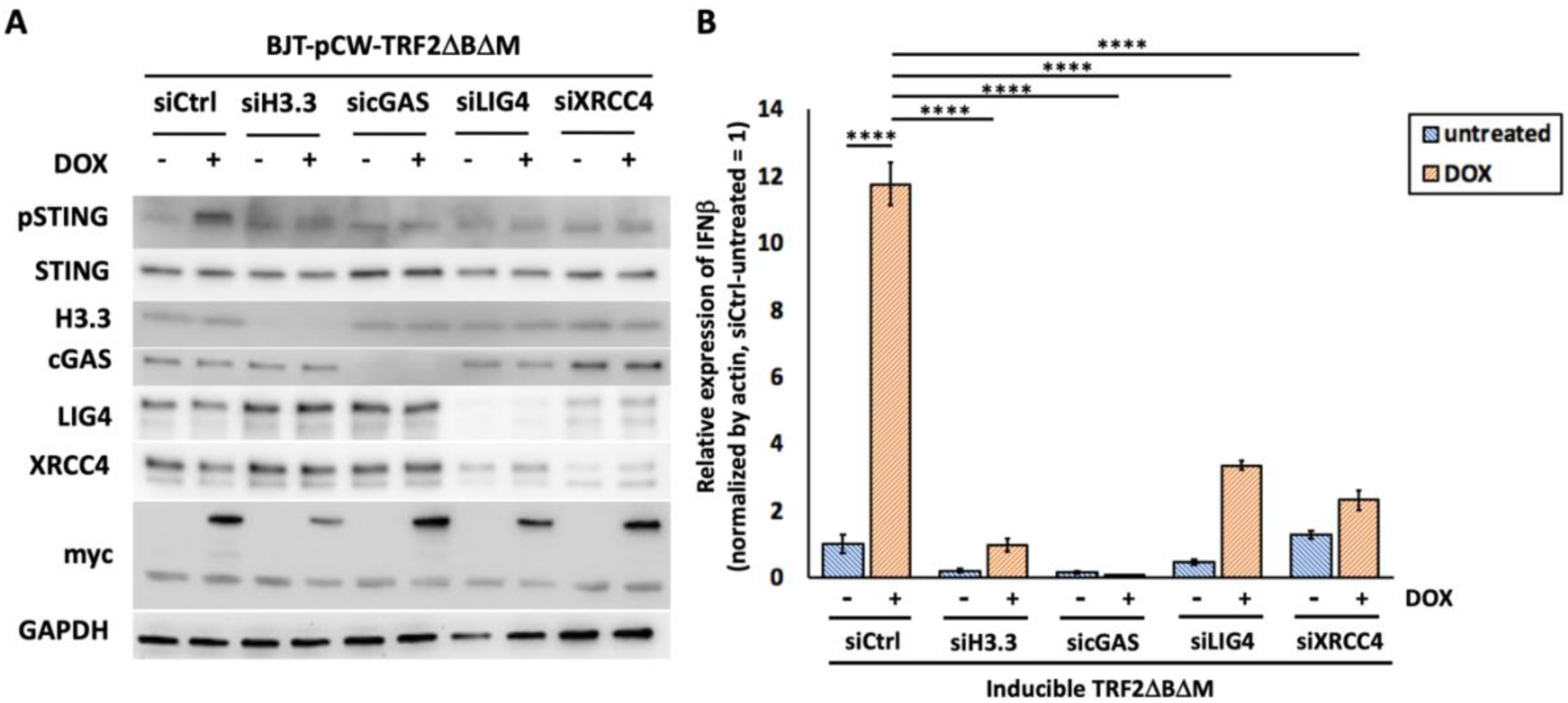
Depletion of LIG4 and XRCC4 disrupts activation of the cGAS-STING response. BJ^t^ cells inducibly expressing myc-tagged TRF2ΔBΔM were treated with siRNAs at 60 h and 12 h before doxycycline (DOX) treatment. To correlate telomere fusion events and nuclear abnormalities, treated cells were lysed on day 2 after DOX treatment. (**A**) Protein levels of phosphorylated STING (pSTING), STING, H3.3, cGAS, LIG4, XRCC4, myc, and GAPDH were determined by Western blotting. (**B**) mRNA expression levels of IFNβ were determined by RT-qPCR. qPCR experiments represent the mean ±SD of three technical repeats. Statistical analysis was conducted using one-way ANOVA. **** p-value < 0.0001.

### Histone H3.3 mediates DDR and telomere fusion for cGAS-STING pathway activation

As shown above, histone H3.3 plays a role in activating the cGAS-STING pathway in BJ^t^ cells with deprotected telomeres (Fig. 2). We hypothesized that histone H3.3 contributes to telomere fusion, which in turn influences cGAS-STING pathway activation. To examine this possibility, we silenced histone H3.3 expression in BJ^t^ cells by siRNAs and analyzed subsequent telomere fusion and nuclear structural abnormalities following TRF2ΔBΔM induction for two days. Our results demonstrate that depletion of histone H3.3 significantly reduced the number of telomere fusion events and partially reduced the formation of micronuclei, nuclear buds, and nucleoplasmic bridges typically induced by TRF2ΔBΔM expression (Fig. 3). These findings indicate that histone H3.3 is required for NHEJ-mediated telomere fusion following telomere deprotection. Our results reveal that chromosomal and nuclear abnormalities occur on the second day after inducting TRF2 mutant expression. Notably, this precedes the peak cGAS-STING response, which occurs on the third day post-induction of TRF2ΔBΔM expression (Fig. 3, S1). These observations suggest that histone H3.3 orchestrates the downstream response to TRF2ΔBΔM-induced telomere deprotection, including telomere fusion, nuclear abnormalities, and subsequent cGAS-STING pathway activation. Since depletion of cGAS did not rescue TRF2ΔBΔM-mediated telomere fusion or nuclear abnormalities (Fig. 3), cGAS does not appear to directly participate in DNA repair at deprotected telomeres but rather functions downstream of telomere fusion and nuclear abnormalities. Thus, our results suggest that histone H3.3 mediates cGAS-STING activation by promoting NHEJ-mediated repair of deprotected telomeres.

Next, we investigated if histone H3.3 regulates the telomere DDR and thereby controls NHEJ for subsequent cGAS-STING activation. As evidenced above, expression of TRF2ΔBΔM elicited moderate TIF formation (Fig. 1B, 5A-C), indicating that telomere deprotection can trigger a DDR. To determine whether histone H3.3 regulates the telomeric DDR, we treated BJ^t^ cells with siRNAs to deplete the expression of histone H3.3. Interestingly, we observed that depletion of histone H3.3 also reduced TIF formation in the TRF2ΔBΔM-expressing cells (Fig. 5A-C), supporting that the role of histone H3.3 in regulating the DDR at deprotected telomeres. These findings underscore the role of histone H3.3 in regulating the TRF2ΔBΔM-induced telomeric DDR.

**Figure 5.**
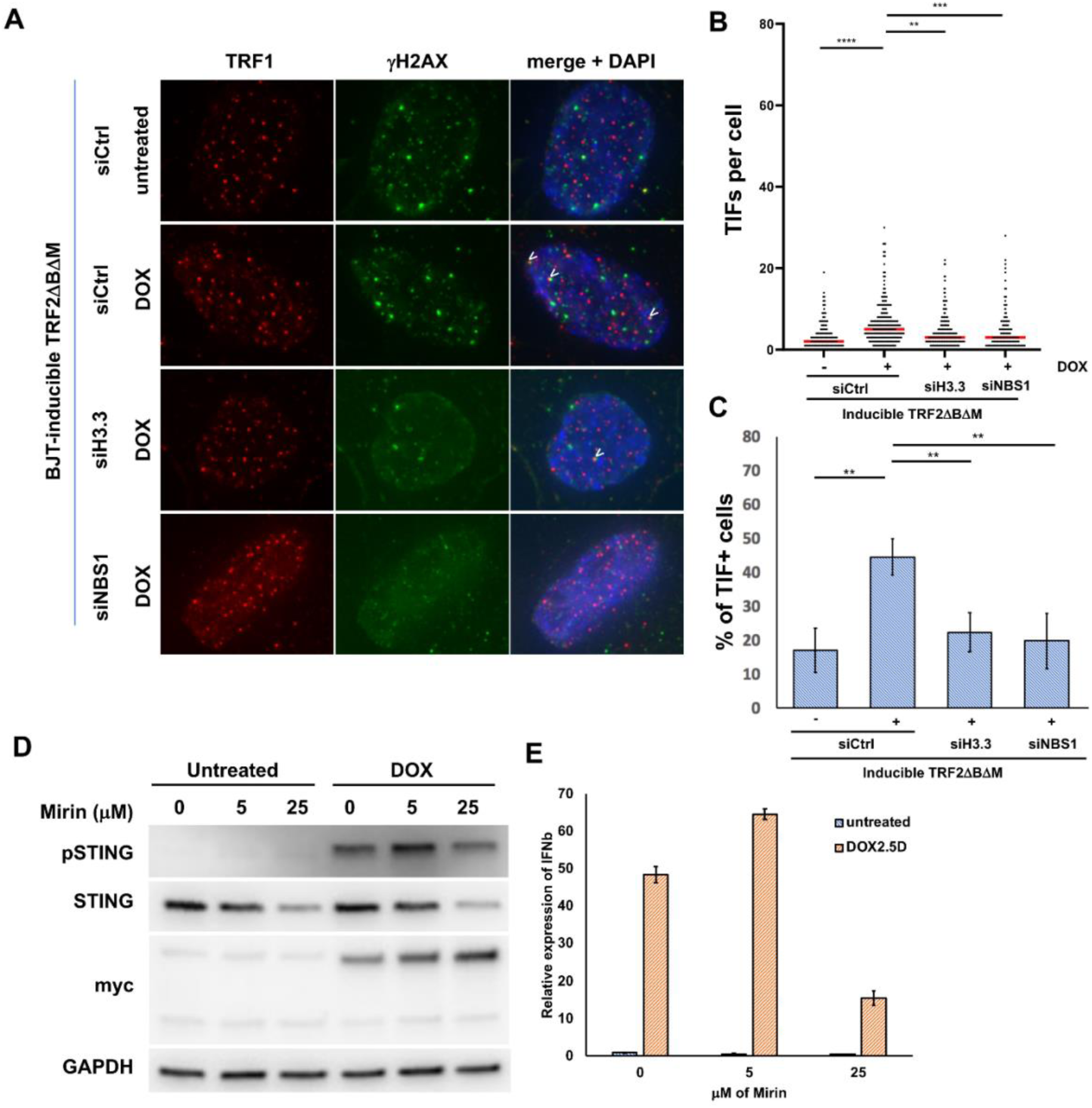
Depletion of H3.3 expression limits TIF formation. BJ^t^ cells were treated with siRNAs including siCtrl, siH3.3, and siNBS1 at 60 h and 12 h before inducing stable expression of TRF2ΔBΔM or TPP1ΔRD through 2.5 ng/ml DOX treatment. (**A**) Immunostaining for γH2AX and TRF1 was performed on day 2 after DOX treatment. Representative images stained for γH2AX (green) and TRF1 (red). White arrowheads indicate representative colocalized signals of TRF1 and γH2AX, which are defined as TIFs, in the cells. (**B**) Numbers of TIFs per cell for the different treatment groups. Each dot represents the TIF count per cell. Red lines indicate the median value for each group. The data represent three biological repeats, with 70-120 cells counted in each repeat, and each experimental group represents 250-330 cells. (**C**) Percentages of cells harboring >5 TIFs. More than 100 cells were quantified for each sample in each repeat. BJ^t^ cells inducibly expressing either TRF2ΔBΔM were treated with 0, 5, or 25 μM of MRN complex inhibitor, Mirin. Total RNA and protein samples are collected after doxycycline treatment for 2.5 days. (A) Protein levels of phosphorylated STING (pSTING), STING, myc, and GAPDH were determined by Western blotting. (B) Relative expression levels of IFNβ mRNA in each sample group were determined by qPCR. qPCR experiments represent the mean ±SD of three technical repeats. Data represent mean ±SD of three biological repeats. Statistical analysis was conducted using one-way ANOVA. ** p-value < 0.01, *** p-value <0.001, and **** p-value < 0.0001.

Since NBS1 is a key component of the MRN complex, we hypothesized that its depletion would impair telomeric DDR. The results showed that TRF2ΔBΔM-induced TIF formation was abolished upon NBS1 depletion (Fig. 5A-C), consistent with the MRN complex playing a role in sensing telomere deprotection to activate a DDR^46^. To investigate whether the MRN complex regulates cGAS-STING activation, we measured IFNβ production in cells treated with the MRN inhibitor, Mirin^47^. Treatment with 25 μM Mirin significantly reduced STING phosphorylation and IFNβ transcription in TRF2ΔBΔM-expressing cells, confirming that MRN is essential for cGAS-STING activation following telomere deprotection (Fig. 5D, E). Our observations further demonstrate that the cellular machinery responsible for telomeric DDR also modulates the activation of cGAS-STING pathway. Collectively, these results indicate that histone H3.3 and MRN complex facilitate DDR activation and NHEJ at deprotected telomeres, thereby priming cells for subsequent cGAS-STING pathway activation.

### Histone H3.3’s chaperones are required for TRF2ΔBΔM-driven cGAS-STING pathway activation, telomere fusion, and TIF formation

Histone chaperones, including ATRX-DAXX and HIRA, have been shown to mediate the deposition of histone H3.3 on telomeres^18,48^. Anti-silencing function 1 (ASF1) binds histone H3.3-H4 dimers and transfers them to HIRA^48^. To investigate if deposition of histone H3.3 on telomeres is important for regulating telomere deprotection-induced cGAS-STING activation, we used siRNAs to deplete individual histone H3.3 chaperone proteins, including ASF1, HIRA, ATRX, and DAXX, from BJ^t^ cells expressing TRF2ΔBΔM. Consistently, depletion of ATRX, DAXX, or HIRA reduced the telomeric distribution of histone H3.3, as evidence by decreased telomeric signals in histone H3.3 chromatin immunoprecipitation (ChIP) assays following siRNA-mediated depletion of ATRX, DAXX, or HIRA siRNAs (Fig. S2). STING phosphorylation and IFNβ expression were reduced upon TRF2ΔBΔM expression in the absence of ATRX or DAXX (Fig. 6A, 6C). Additionally, depletion of HIRA or ASF1 significantly reduced STING phosphorylation and IFNβ expression in TRF2ΔBΔM-expressing cells (Fig. 6B, 6C). Together, these observations indicated that ATRX, DAXX, HIRA, and ASF1 are all required to activate the cGAS-STING response caused by telomere deprotection.

**Figure 6.**
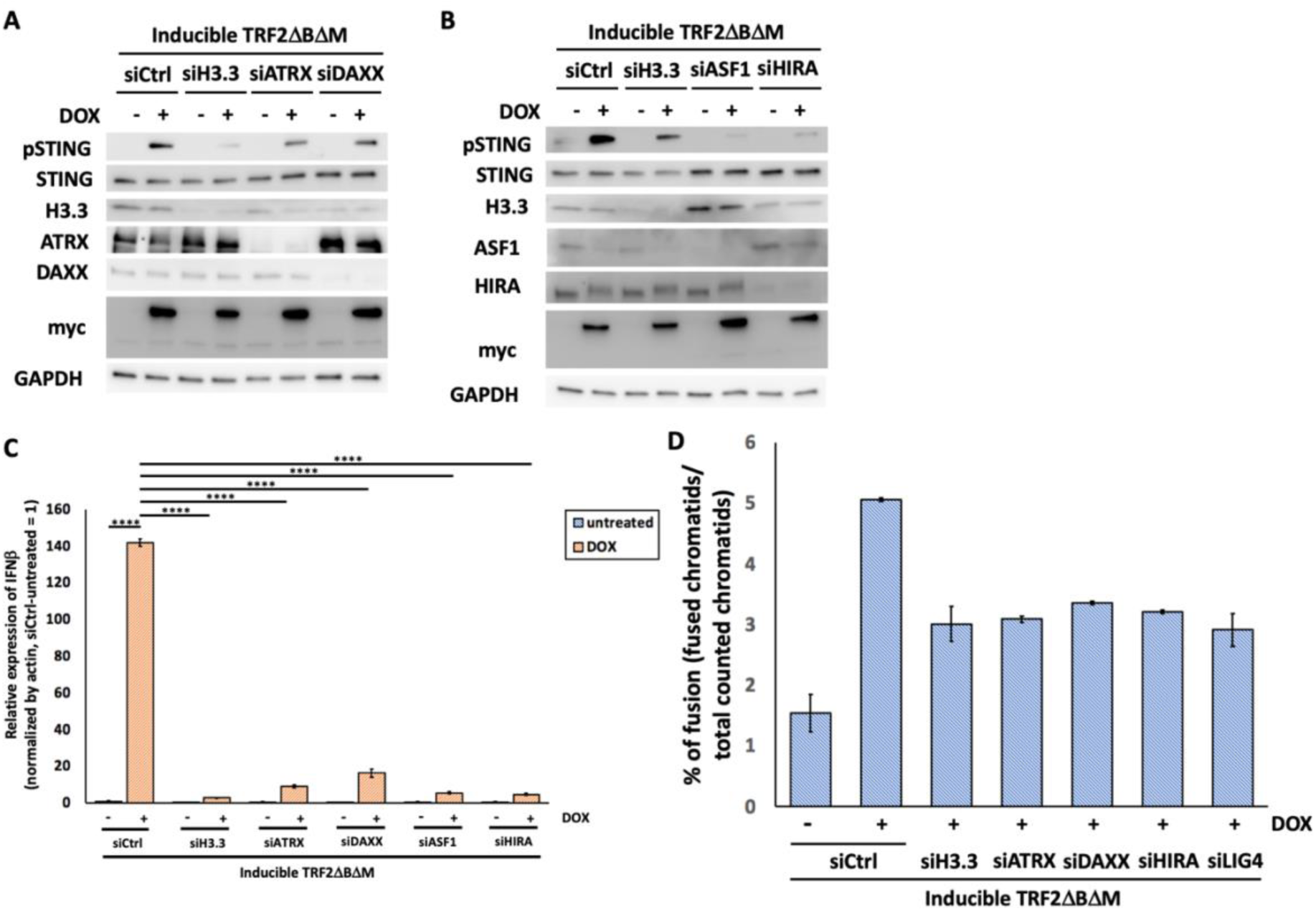
Depletion of H3.3 chaperones impairs cGAS-STING pathway activation. BJ^t^ cells were treated with siRNAs at 60 h and 12 h before inducing expression of myc-tagged TRF2ΔBΔM through DOX treatment. Samples were cultured for 2.5 days to investigate the cGAS-STING signaling response. (**A**, **B**) Protein levels of phosphorylated STING (pSTING), STING, ASF1, HIRA, ATRX, DAXX, H3.3, myc, and GAPDH were determined by Western blotting. (**C**) mRNA expression levels of IFNβ were determined by RT-qPCR. qPCR experiments represent the mean ±SD of three technical repeats. Another set of BJ^t^ cells was treated with siRNAs including siCtrl, siH3.3, siATRX, siDAXX, siHIRA, and siLIG4 at 60 h and 12 h before DOX treatment to induce expression of TRF2ΔBΔM for 24 h. Cells were arrested using 0.1 μg/ml Demecolcine for another 24 h, and then spread on a slide to visualize telomere fusion events by staining with a TelC probe. Statistical analysis was conducted using one-way ANOVA. **** p-value < 0.0001. (**D**) Quantitative results for telomere fusion events were calculated from >2000 chromosomes for each sample. Data represent the mean ±SD of two sets of experiments.

To determine how these histone chaperones regulate downstream responses to telomere deprotection, we assessed telomeric DDR and telomere fusion in cells depleted of ATRX, DAXX, or HIRA. Depletion of ATRX, DAXX, or HIRA reduced the percentages of fused chromatids and TIF formation in TRF2ΔBΔM-expressing BJ^t^ cells (Fig. 6D, S3), indicating that proper telomeric distribution of histone H3.3 is essential for DDR activation and telomere fusion following telomere deprotection. Thus, histone H3.3 localization at telomeres is essential for DDR activation, telomere fusion, and cGAS-STING pathway activation.

## DISCUSSION

Our results reveal a role for histone H3.3 in mediating telomere deprotection-induced activation of the cGAS-STING pathway. Although histone H3.3 deposition at telomeres has been previously documented, its role in telomere dynamics and the DNA damage response remains poorly understood. Notably, mutations in *H3F3A*, the gene encoding histone H3.3, are strongly associated with glioblastoma, a tumor that predominantly relies on the ALT mechanism^28,49^. Our previous work demonstrated that ALT cancer cells lack a functional cGAS-STING-mediated intracellular DNA sensing mechanism and that histone H3.3 and its chaperone contribute to regulating this pathway^15^. Here, we show that histone H3.3 contributes to telomere fusion events and the formation of TIFs in cells with deprotected telomeres (Fig. 3, 5). These findings suggest that histone H3.3 contributes to ALT cancer-associated telomere dysfunctions, including TIF formation, chromosome fusion events, and cGAS-STING pathway activation.

Recent studies have shown that glioblastoma-associated histone H3.3 variants, such as histone H3.3^K27M^ and histone H3.3^G34V^, impair chromosome segregation, trigger genetic instability^25^, and regulate ALT-associated features^50^. These findings suggest that ALT-associated histone H3.3 mutations may suppress the DDR, promote genetic instability, and facilitate error-prone genome doubling and rapid cell cycle progression. These mutations may also suppress cGAS-STING activation, dampening the innate immune response and allowing cancer cells to thrive under genomic stress. However, further studies are needed to clarify the relationship between histone H3.3 mutations and cGAS-STING pathway activity in ALT cancer cells to better understand how these mutations contribute to ALT phenotypes.

ALT cancers maintain telomeres through a telomerase-independent recombination-driven process. Given the link between histone H3.3 and ALT cancers^25–28^, we propose that its expression and telomeric deposition influence DNA repair pathway choice, shifting the balance between NHEJ and homologous recombination (HR). Histone H3.3 has been shown to function downstream of the PARP1-CHD2 machinery to promote NHEJ at telomeres^37^. Similarly, the histone H3.3’s chaperone molecule, ATRX is active in both NHEJ^33^ and HR^51^. In our observations, depletion of histone H3.3 or its chaperones (ATRX, DAXX, and HIRA) reduces telomere fusion events (Fig. 3, 6) and impairs TIF formation (Fig. 5, S3), underscoring their importance in telomere repair. Since ALT phenotypes depend on HR, we propose that mutations in histone H3.3 or its chaperones disrupt NHEJ while enhancing HR, thereby supporting ALT telomere maintenance. It will be important to establish in the future how histone H3.3 localization and chaperone interactions mediate this switch.

Telomere fusion has been proposed as a mechanism to counteract telomere shortening^52,53^. However, our findings reveal that telomere fusion also activates the cGAS-STING pathway, which may impair cell growth. This scenario is evidenced by the impairment of cGAS-STING activation we detected upon depletion of the DNA repair proteins LIG4 and XRCC4 (Fig 4). Furthermore, cGAS depletion rescues the growth inhibition induced by telomere deprotection (Fig. 1E, F), suggesting that telomere fusion promotes cellular senescence via cGAS-STING pathway activation. Interestingly, our findings indicate that cGAS-STING activation occurs in a pulsatile manner following telomere deprotection (Fig. S1). Thus, telomere fusion appears to trigger rapid but transient activation of this pathway under telomeric stress. Notably, transient activation of the cGAS-STING pathway by TRF2ΔBΔM expression is sufficient to inhibit cell growth (Fig. 1E, F). These findings reveal a novel role for telomere fusion in transient cGAS-STING pathway activation and cell growth regulation. Future studies should explore the mechanisms underlying this temporal control and its broader implications. Overall, our results highlight the essential role of histone H3.3 in deprotected telomere-driven cGAS-STING signaling and provide new insights into its underlying mechanisms.

## RESOURCE AVAILABILITY

### Lead contact

Requests for further information and resources should be directed to and will be fulfilled by the lead contact, Liuh-Yow Chen.

### Materials availability

This study did not generate new unique reagents.

### Data and code availability

Original Western blot images and microscopy data reported in this paper will be shared by the lead contact. Any additional information required to reanalyze the data reported in this paper is available from the lead contact upon request.

## ACKNOWLEDGEMENT

We thank the Imaging Core of the Institute of Molecular Biology (IMB), Academia Sinica, for technical support. We also thank Nithila Joseph in our lab and John O’Brien for assistance with English editing. This project is supported by grants from Academia Sinica (AS-GCP-113-L02) and the Ministry of Science and Technology (111-2311-B-001-013).

## AUTHOR CONTRIBUTIONS

Conceptualization, C.-M. H. and L.-Y. C.; methodology, C.-M. H.; analysis, C.-M. H.; investigation, C.-M. H.; resources, L.-Y. C.; writing – original draft, C.-M. H. and L.-Y. C.; writing – review and editing, C.-M. H. and L.-Y. C.; funding acquisition, L.-Y. C.; supervision, C.-M. H. and L.-Y. C.

## DECLARATION OF INTERESTS

The authors declare no competing interests.

---

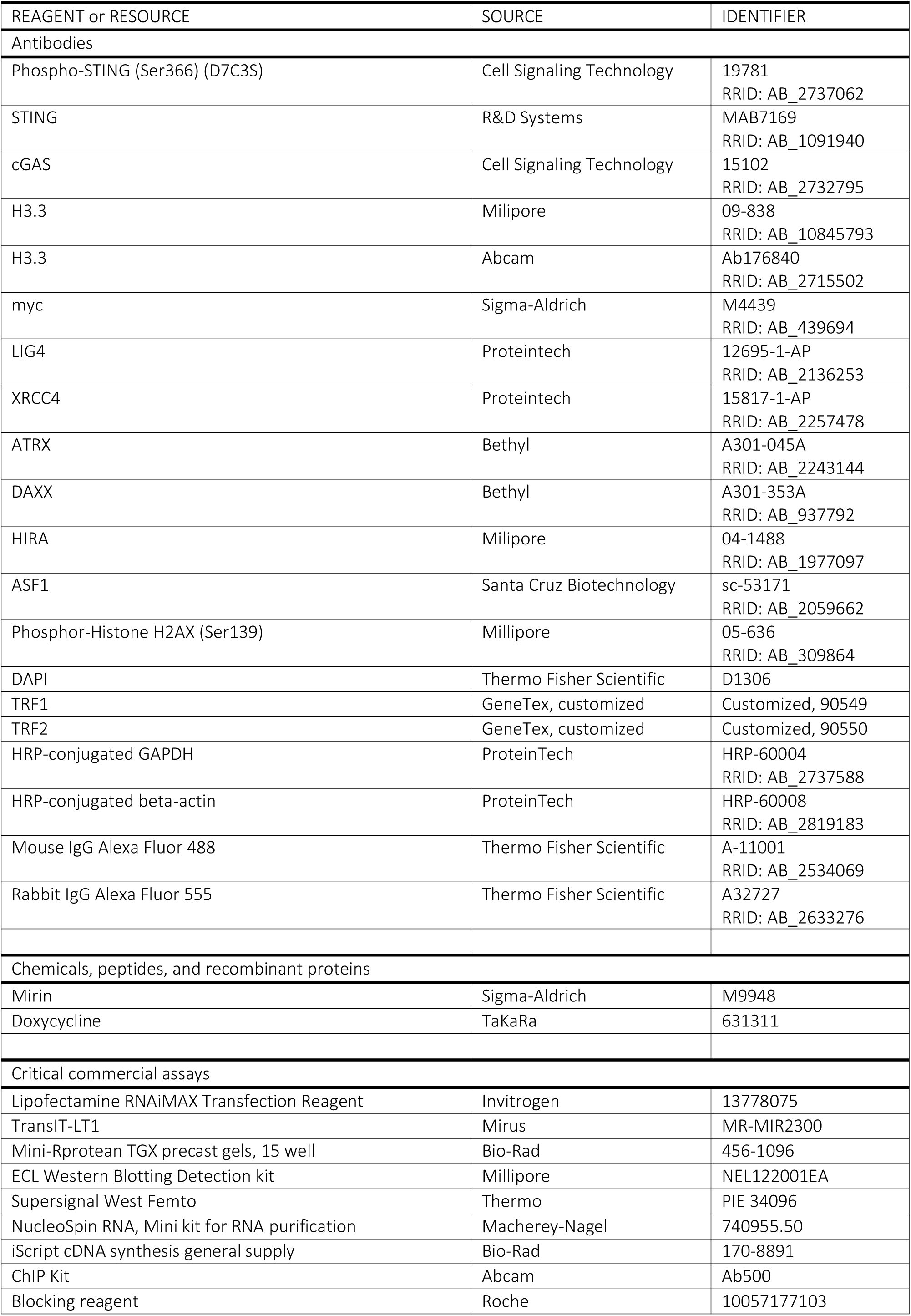
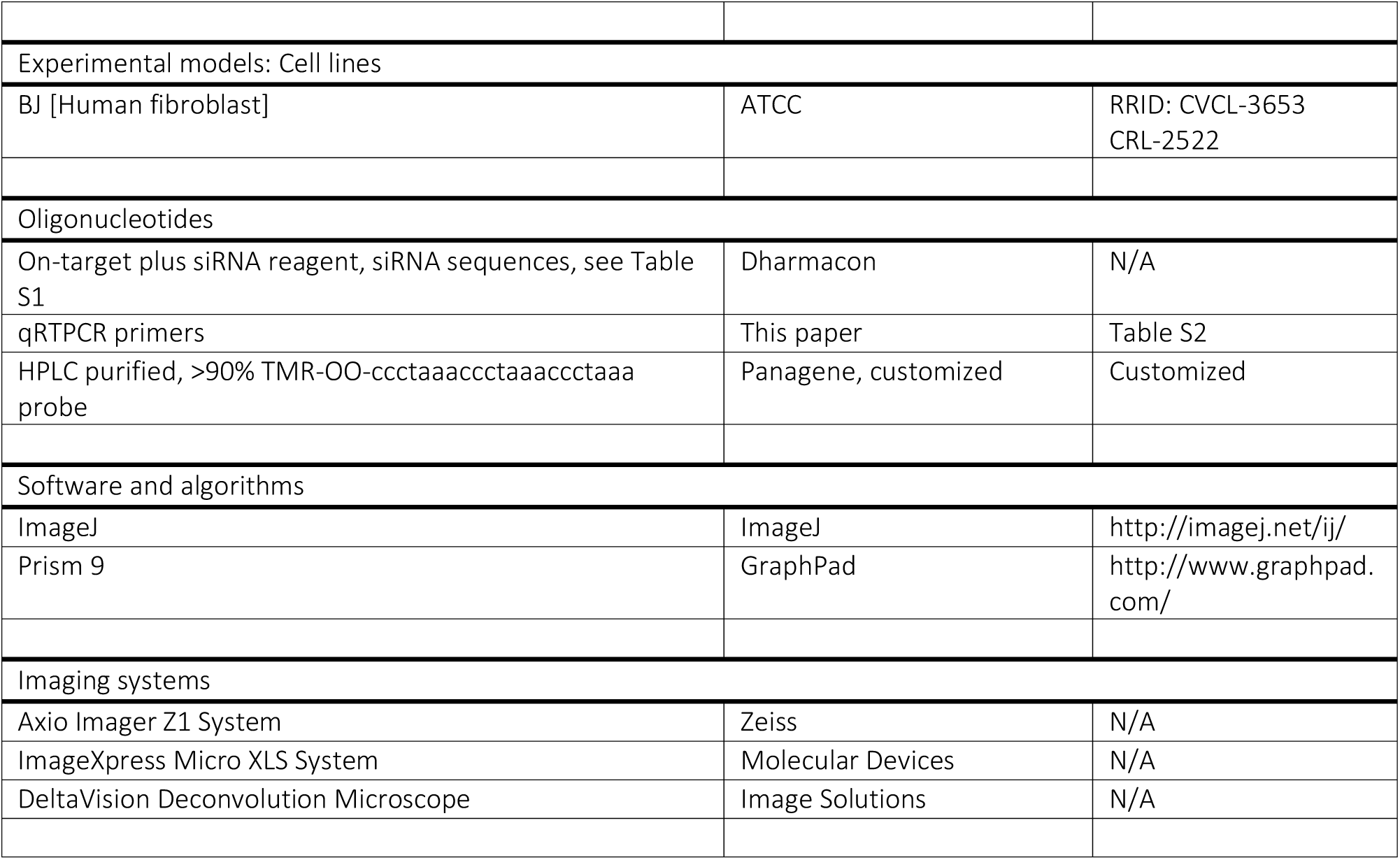

## Materials and Methods

### Cell lines

Cell lines were obtained from ATCC and maintained under their recommended culturing conditions. BJ^t^ cells originated from BJ normal fibroblasts that stably express hTERT. The retroviral transformation was performed to generate cells that inducibly express green fluorescence protein (GFP), the TRF2 mutant TRF2ΔBΔM, or the TPP1 mutant TPP1ΔRD. Cells were tested for target protein expression by treating with a minimum of 25 ng/ml of doxycycline.

### Gene silencing

The siRNAs used were On-target plus siRNA reagents (Dharmacon). The siRNA reagents were generated using the SMARTselection algorithm, with chemical modifications to eliminate off-target effects. Four siRNAs for each of H3F3A, H3F3B, cGAS, LIG4, XRCC4, ATRX, DAXX, HIRA, ASF1A, and ASF1B, as well as two siRNAs for NBS1, were employed. siRNAs were transfected into the cells using Lipofectamine RNAiMAX (Invitrogen) according to the manufacturer’s instructions. Depletion of H3.3 and ASF1 was performed by mixing equal amounts of siRNAs targeting H3.3 and ASF1. MRN chemical inhibitors, Mirin (M9948, Sigma), was dissolved in DMSO as 10mM stock, and treated to the cell culture at the indicated concentrations.

### Protein extraction, Western blots, and antibodies

Cell extracts were harvested using 1 X SDS sample buffer (62.5 mM Tris, pH 6.8, 10% glycerol, 2% SDS, 0.01% bromophenol blue, and 10% β-mercaptoethanol). Proteins were denatured by heating the extract at 95 °C for 10 minutes. Proteins were separated by SDS-PAGE, and transferred to a nitrocellulose membrane. The membranes were blocked with either Tris-buffered saline (TBS) at pH 8.0 with 5% milk and 0.1% Tween-20 to determine phosphorylated proteins or using Phosphate-buffered saline (PBS) at pH 8.0 with 5% milk and 0.1% Tween-20 to determine other proteins. Primary antibodies including phosphorylated STING (19781, Cell Signaling), STING (MAB7169, R&D Systems), cGAS (15102, Cell Signaling), H3.3 (09-838, Merkmillpore; ab176840, Abcam), myc (M4439, Sigma), LIG4 (12695-1-AP, Proteintech), XRCC4 (15817-1-AP, Proteintech), ATRX (A301-045A, Bethyl), DAXX (A301-353A, Bethyl), HIRA (04-1488, Millipore), and ASF1 (sc-53171, Santa Cruz Biotechnology) were used at 1:1000 dilution. Horseradish peroxidase-conjugated secondary antibodies were used at 1:2000 dilution to determine the proteins in the samples. As an internal control for gene expression, HRP-conjugated GAPDH antibody (HRP60004, ProteinTech) and HRP-labeled ACTB (HRP-60008, ProteinTech) were used at 1:5000 dilution. Either enhanced chemiluminescence (ECL) or SuperSignal West Femto Maximum Sensitivity reagents (Thermo Fisher Scientific) were used for signal detection.

### RNA extraction and quantitative real-time PCR

Cells were lysed and harvested to extract total RNA using an RNA Mini Kit (Macherey-Nagel). Reverse transcription was performed with iScript Reverse Transcriptase Supermix (Bio-Rad) to generate cDNA for quantitative analyses. Quantitative real-time PCR was performed using SYBR Green Select reagent (LifeTechnologies) to determine expression levels of target genes. Quantitative PCR analysis was performed using a QuantStudio 12K Flex Real-Time PCR System (ThermoFisher Scientific). The primer sequences are as follows: β-actin, 5’-AGCACTGTGTTGGCGTACAG-3’ and 5’-TCCCTGGAGAAGAGCTACGA-3’; interferon β, 5’-AAACTCATGAGCAGTCTGCA-3’ and 5’-AGGAGATCTTCAGTTTCGGAGG-3’. Data analysis was performed by calculating the comparative Ct (ΔΔCt) of target genes normalized to an internal control (β-actin).

### Metaphase spread and telomere FISH on chromosomes

Analysis of telomere-to-telomere fusion was performed in DOX-treated cells subjected to siRNA-mediated gene expression depletion. Cells were treated with DOX for 30 hours, and reseeded in 0.1 μg/ml Demecolcine-containing medium. After 18 h of incubation, the floating and attached cells were collected for further analysis. Cells were incubated with 75 mM KCl for 15 mins at 37 °C and routinely mixed at 5-minute intervals. Following KCl incubation, the cells were fixed with fixing buffer (3:1 methanol: acetic acid) at 4 °C for more than 24 h. Fixed cells were dropped on a glass slide from a height of 20 cm, heated at 75 °C for 1 min, and then dried at room temperature overnight. The slides were stained according to a standard fluorescence in situ hybridization (FISH) FISH protocol. In brief, the slides were fixed with 10% formalin for 2 mins, and then dehydrated by incubating them in increasing percentages of ethanol solution. The slides were dried and incubated with a probe mixture (70% formamide, 0.5% blocking reagent (Roche), 250 nM TMR-conjugated (CCCTAA)3-PNA probe, 10 mM Tris pH 7.4) at 80 °C for 3 mins. The slides were hybridized at room temperature in the dark for 3 h. Serial wash steps were then performed, including twice with Wash I (70% formamide, 10 mM Tris-HCl pH 7.5), and twice with Wash II (100 mM Tris-HCl pH 7.5, 150 mM NaCl, 0.08% Tween-20). The slides were counterstained with DAPI, dehydrated with ethanol, and mounted with Fluoromount Aqueous Mounting Medium (Sigma). Images were acquired with a Zeiss Axio Imager Z1 System. Determination and quantitative analysis of telomere fusion events were performed by counting for 700 to 1500 chromosome pairs.

### Immunofluorescence staining

Cells grown on coverslips were treated with permeabilization solution (20 mM Hepes pH 8.0, 50 mM NaCl, 3 mM MgCl2, 0.5% Triton-X 100, 300 mM sucrose), and fixed with 2% paraformaldehyde. The cells were permeabilized with 0.5% Triton-X 100, and then blocked with 2% fetal bovine serum (FBS) in phosphate-buffered saline. Primary antibodies including anti-TRF1 (90549, GeneTex) and anti-γH2AX (05-636, Millipore) were used. The secondary antibodies were Mouse IgG Alexa Fluor 488 (A-11001, Thermo Fisher Scientific) and Rabbit IgG Alexa Fluor 555 (A32727, Thermo Fisher Scientific). The cells were stained with DAPI, dehydrated, and mounted with ProLong Gold Antifade Mountant (Life Technologies). Images were acquired under a DeltaVision Deconvolution Microscope. Colocalization events were determined based on the overlapping of signals from different channels, and quantification was performed by counting for > 100 nuclei on one coverslip.

### Chromatin immunoprecipitation (ChIP)

The chromatin immunoprecipitation protocol was adapted from the Abcam ChIP kit (Ab500). After depletion of gene expression by siRNAs for 4 days, the BJ^t^ cells were fixed with 1% formaldehyde for 10 min. The cross-linking process was quenched with glycine, and the cells were collected from the culture dish. Chromatin DNA was extracted from cells, and sheared into 200-500 bp. Sheared chromatin DNAs were immunoprecipitated with antibodies, including H3.3 (09-838, Merkmillpore) and TRF2 (90550, GeneTex) antibodies. Antibody-precipitated chromatin DNAs were enriched with protein A beads. The enriched chromatin DNAs were reversed crosslinking with protease and extracted from the precipitated DNA pellet. Precipitated DNAs were then loaded on the Hybond-XL membrane. The telomeric DNA from each sample was determined with α-P32-dCTP-labelled telomere probe.

### Cell growth and quantification of nuclear abnormalities

Cell growth was determined by counting cells in the multi-well culture plates. Cells were plated in 12-well plates and fixed with 2% paraformaldehyde on the 1^st^-7^th^ days post-treatment. The cells were stained with DAPI and imaged using an ImageXpress Micro XLS System (Molecular Devices). For each well, images in 64 fixed regions were taken to quantify the number of cells inside the well. These images were used to quantify nuclear abnormalities, including nucleoplasmic bridges, nuclear buds, and micronuclei. Such abnormalities were identified based on a deformed nuclear structure, and the quantification was performed by counting for >2000 nuclei in one sample.

### Quantification and statistical analysis

With the help of GraphPad Prism, a one-way analysis of variance (ANOVA) was performed to assess the statistical differences among group means. Following a significant ANOVA result (P<0.05), post-hoc comparisons were conducted using Tukey’s Honest Significant Difference (HSD) test to determine pairwise difference between group means and to control for Type I error due to multiple comparisons.

## Notes

### Competing Interest Statement

The authors have declared no competing interest.

### Summary of Updates

Updated the abstract, introduction, and discussion sections. Paragraph wording clarified

