## Supplemental figures and tables for "Histone Variant H3.3 Mediates DNA Repair and cGAS-STING Pathway Activation in Telomere Dysfunction"

### Supplementary Data

**Figure S1.**

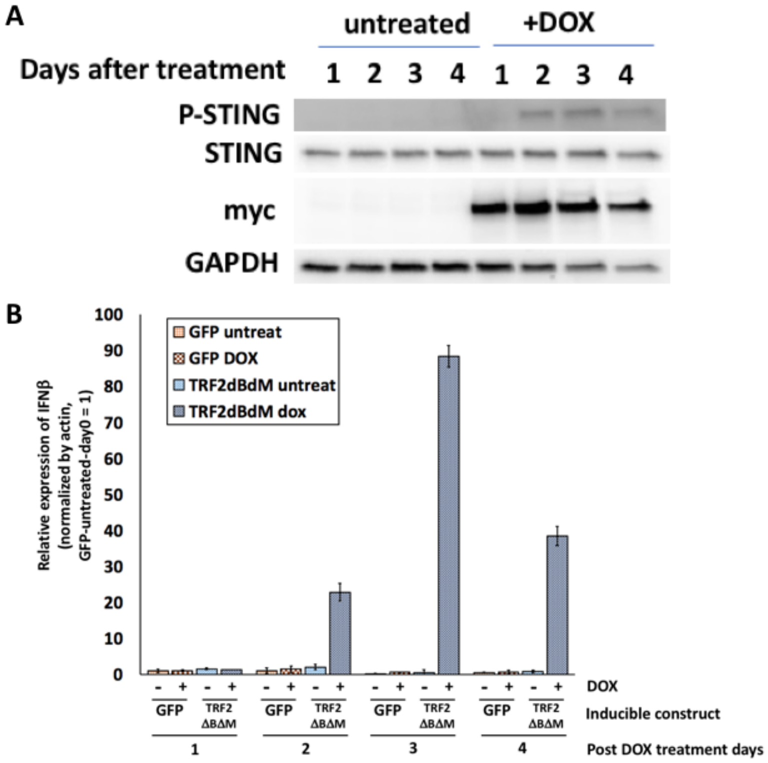


Expression of TRF2ΔBΔM triggers a cGAS-STING response within 4 days of inducing expression of TRF2ΔBΔM. BJ$^{t}$ cells inducibly expressing either GFP or TRF2ΔBΔM were treated with 25 ng/ml doxycycline (DOX) for 1 to 4 days before being harvested for protein or RNA. (**A**) Protein levels of phosphorylated STING (pSTING), STING, myc, and GAPDH were determined by Western blotting. (**B**) Relative expression levels of IFNβ mRNA in each sample group were determined by qPCR. qPCR experiments represent the mean ±SD of three technical repeats.

**Figure S2.**

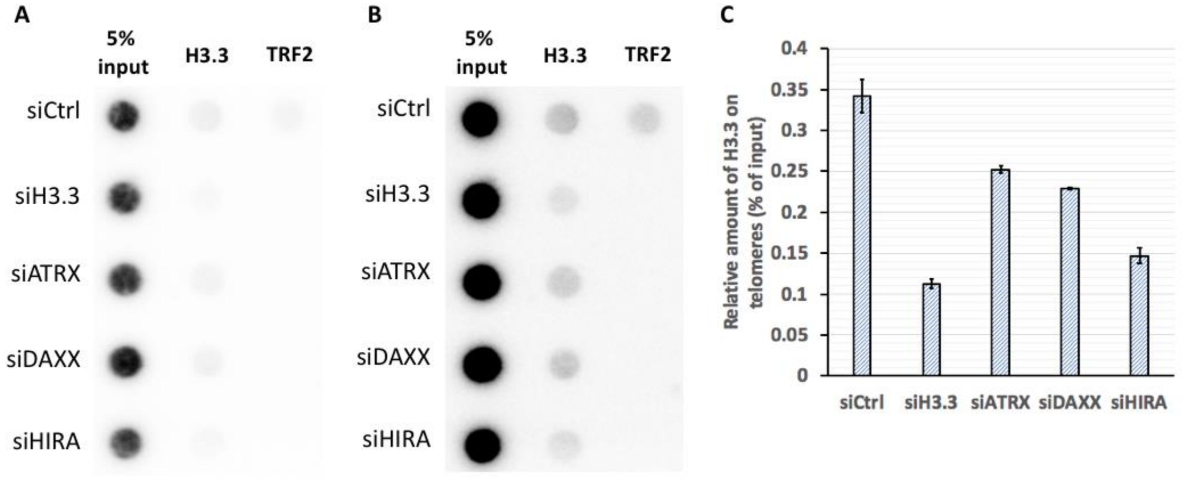


Depletion of ATRX, DAXX, and HIRA all contributes to a decreased amount of H3.3 on telomeres. Chromatin DNA was harvested from $BJ^T$ cells treated with siRNAs, including siCtrl, siH3.3, siATRX, and siHIRA, for 4 days. The DNA samples were sheared into 200-500 bp, and pulled down with H3.3 or TRF2 antibodies. 5% of input DNA and antibody-precipitated DNA samples were isolated from DNA-binding proteins, and loaded on a Hybond-XL paper for blotting. Radioactive isotope-labeled telomeric probes were used to show the telomere signals in each sample. (**A**) Dot-blot results for telomeric signals in samples. (**B**) enhanced contrast result of (**A**). (**C**) Relative amount of H3.3 on telomeres. Data represent the mean ±SD of technically duplicated results.

**Figure S3.**

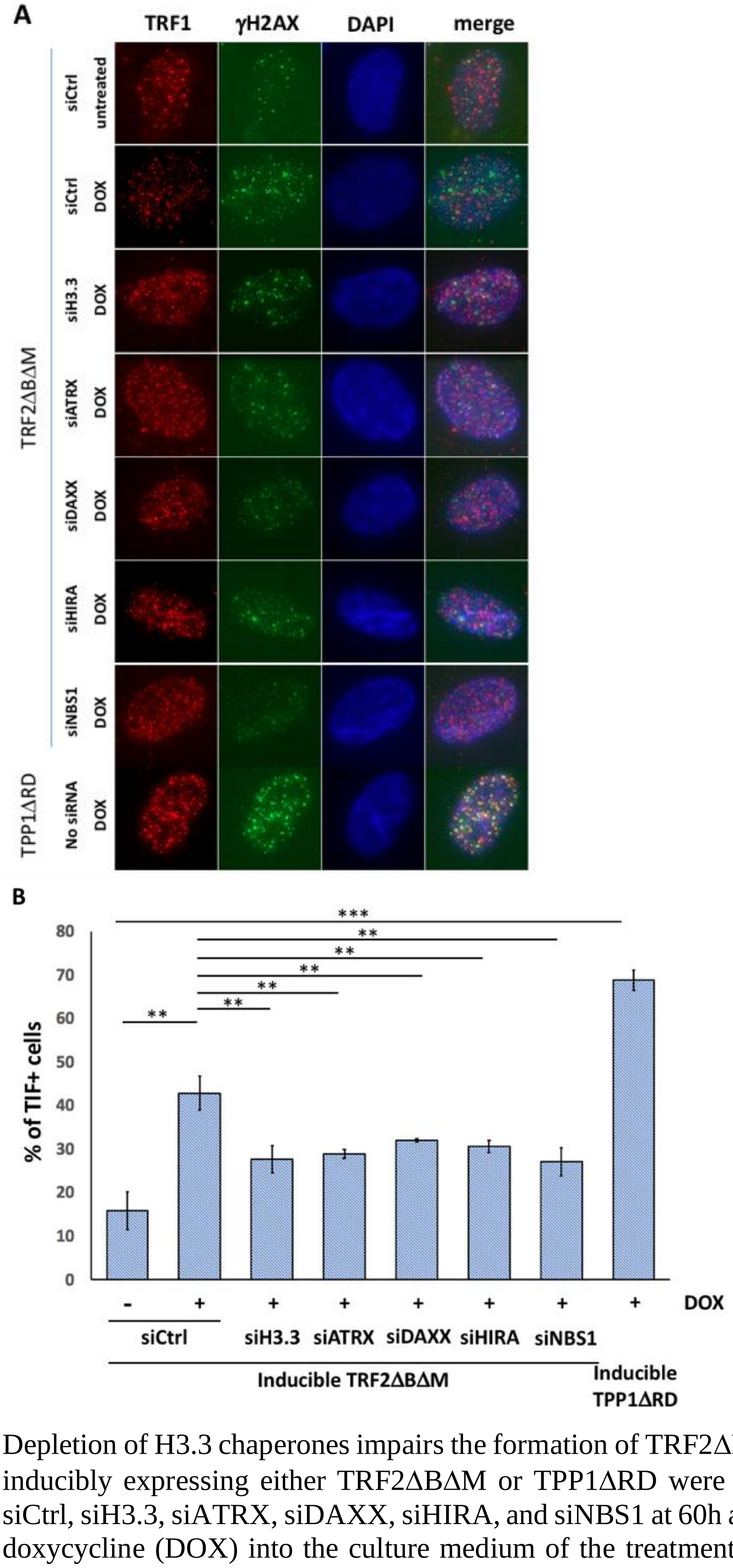


Depletion of H3.3 chaperones impairs the formation of TRF2ΔBΔM-dependent TIFs. BJ$^{T}$ cells inducibly expressing either TRF2ΔBΔM or TPP1ΔRD were treated with siRNAs including siCtrl, siH3.3, siATRX, siDAXX, siHIRA, and siNBS1 at 60h and 12 h before adding 25 ng/ml doxycycline (DOX) into the culture medium of the treatment group to induce expression of TRF2ΔBΔM or TPP1ΔRD. After 48 h of DOX treatment, the cells were harvested for antibody staining. Cells were stained with anti-TRF1 antibody to indicate telomeres and anti-γH2AX to illuminate TIFs. (**A**) Representative images stained for γH2AX (green) and TRF1 (red). (**B**) Percentages of cells with >5 TIFs. For each sample in each repeat, > 100 cells were quantified. Data represent the mean ±SD of three biological repeats.

**Table S1**

| Targer | Sequences |
|---|---|
| none-targeting | UGGUUUACAUGUCGACUAA |
| none-targeting | UGGUUUACAUGUUGUGUGA |
| none-targeting | UGGUUUACAUGUUUUCUGA |
| none-targeting | UGGUUUACAUGUUUUCCUA |
| H3F3A | CCUAAGUAUAUGAUUGCGA |
| H3F3A | AUAUAAAUGCGGAGACGUA |
| H3F3A | GCAACUAAAUGGUGUUUGU |
| H3F3A | UUUCAGCGGUUCAACUUUA |
| H3F3B | UUAAGAUGGUGUUGGGUUA |
| H3F3B | CAUAGAAGGCUCUUAGAUU |
| H3F3B | GCAGAAGGGAUGGGUGAUA |
| H3F3B | GUGCGAUGAUGGCCACUAA |
| MB21D1 (cGAS) | GAAGAAACAUGGCGGCUAU |
| LIG4 | SMARTPool* |
| XRCC4 | SAMRTPool* |
| ATRX | GAUGUUAGCUGGAUGUGUU |
| ATRX | AGUCAUAGAUGCUAAGUUU |
| ATRX | GCUUGAGGUUUCUGAAUUA |
| ATRX | GUACAGGCGUUAGCAUUAA |
| DAXX | CAGCCAAGCUCUAUGUCUA |
| DAXX | GAGGUUAACAGGCGCAUUG |
| DAXX | GCAAAACAAAGGACGCAUA |
| DAXX | GGAGUUGGAUCUCUCAGAA |
| HIRA | CAUGGGACCCUGUUGGUAA |
| HIRA | GGAUAACACUGUCGUCAUC |
| HIRA | GCUCCGAUCCUUCCAUGUA |
| HIRA | GCAGGCGAUUCUGUCAAUA |
| ASF1A | CGAUCAAGUUUUAGACUCU |
| ASF1A | CAUUAGACCAGGUUGUAAA |
| ASF1A | AGCCAUAUGAUGCAGGUUA |
| ASF1A | GGCAUAUGUUUGUAUUUCA |
| ASF1B | GCACUCCUAUCAAGGGCUU |
| ASF1B | GACAGGAGUUCAUCCGAGU |
| ASF1B | CGGACGACCUGGAGUGGAA |
| ASF1B | GCAGGGAGACACAUGUUUG |
| NBN (NBS1) | CCAACUAAAUUGCCAAGUA |
| NBN (NBS1) | CCAACUAAAUUGCCAAGUA |
| NBN (NBS1) | GCAGAUACAUGGGAUUUGA |
| NBN (NBS1) | GCAGAUACAUGGGAUUUGA |

Sequence information of siRNAs for indicated targeted genes. * is the pooled siRNA product from requesting SMARTpool services.

**Table S2**

| Target gene | forward primer sequence | reverse primer sequence |
|---|---|---|
| beta-actin | AGCACTGTGTTGGCGTACAG | TCCCTGGAGAAGAGCTACGA |
| IFN beta | AAACTCATGAGCAGTCTGCA | AGGAGATCTTCAGTTTCGGAGG |

Primer sequence information for RT-PCR.
